## Supplementary Materials for "Mesoscale simulation of biomembranes with FreeDTS"

### Section 1: Discrete shape operator, principal directions, and curvatures

**Vertex normal and area:** On each vertex, a normal ($\hat{\boldsymbol{n}}_v$) and an area associated with the vertex ($A_v$) are defined as

$$A_v = \frac{1}{3}\sum_{T(R)} A_T \qquad (1.1)$$

$$\hat{\boldsymbol{n}}_v = \frac{\sum_{T(R)} A_T \hat{\boldsymbol{n}}_T}{\left|\sum_{T(R)} A_T \hat{\boldsymbol{n}}_T\right|} \qquad (1.2)$$

Where $\sum_{T(R)} \ldots$ indicates that the sum is over faces (triangles) in a ring around the vertex. $A_T$ and $\hat{\boldsymbol{n}}_T$ are the triangle area and normal.

**Edge normal:** A normal associated with an edge ($\hat{\boldsymbol{n}}_e$) is obtained using normals of two triangles sharing the edge ($\hat{\boldsymbol{n}}_{T1}, \hat{\boldsymbol{n}}_{T2}$) as

$$\hat{\boldsymbol{n}}_e = \frac{\hat{\boldsymbol{n}}_{T1} + \hat{\boldsymbol{n}}_{T2}}{|\hat{\boldsymbol{n}}_{T1} + \hat{\boldsymbol{n}}_{T2}|} \qquad (1.3)$$

**Shape operator on an edge:** The shape operator on an edge $\boldsymbol{e}$ is be obtained as

$$\overleftrightarrow{\mathbf{S}}_{\boldsymbol{e}} = H_e(\vec{\boldsymbol{r}}_{\boldsymbol{e}} \times \hat{\boldsymbol{n}}_e) \otimes (\vec{\boldsymbol{r}}_{\boldsymbol{e}} \times \hat{\boldsymbol{n}}_e) \quad (1.4)$$

Where $H_e = 2|\vec{r}_e|\cos\frac{\Phi}{2}$ and $\Phi$ is the signed dihedral angle between the two triangles ($\hat{\boldsymbol{n}}_{T1}$ and $\hat{\boldsymbol{n}}_{T2}$) sharing the edge $e$ and $\vec{r}_e$ is the unit vector along edge $\boldsymbol{e}$.

**Shape operator on a vertex:** The shape operator on each vertex ($\overleftrightarrow{\mathbf{S}}_{\boldsymbol{v}}$) is obtained as

$$\overleftrightarrow{\mathbf{S}}_{\boldsymbol{v}} = \frac{\mathbf{1}}{A_v}\sum_{e(R)} (\hat{\boldsymbol{n}}_v . \hat{\boldsymbol{n}}_e) \overleftrightarrow{\mathbf{P}}_{\boldsymbol{v}}^{\dagger} \overleftrightarrow{\mathbf{S}}_{\boldsymbol{e}} \overleftrightarrow{\mathbf{P}}_{\boldsymbol{v}} \quad (1.5)$$

Where $\sum_{e(R)} \ldots$ indicates that the sum is over edges in a ring around the vertex and $\overleftrightarrow{\mathbf{P}}_{\boldsymbol{v}}$ is a projection operator that projects geometrical objects on the tangent plane of the vertex $v$ that is given by

$$\overleftrightarrow{\mathbf{P}}_{\boldsymbol{v}} = \mathbb{I} - \hat{\boldsymbol{n}}_v \otimes \hat{\boldsymbol{n}}_v \qquad (1.6)$$

**Principal curvatures and directions:** Eigenvector $\overleftrightarrow{\mathbf{S}}_v$ are vertex normal, and principal directions $(\hat{\mathbf{T}}_1, \hat{\mathbf{T}}_2, \hat{\boldsymbol{n}}_v)$ while eigenvalues are $c_1$, $c_2$ and zero $(c_1, c_2, 0)$.

### Section 2: Constant tension algorithm (dynamic box)

To simulate a membrane with constant frame tension ($\tau$), there is an extra move to change the membrane frame area. Let's assume our framed membrane has a projected area $A_p$. on $XY$ plane in a rectangular box ($Lx \quad Ly \quad Lz$) with periodic boundary conditions in all three directions. To each Monte Carlo Sweep (MCS) with a probability of $P = 1/(N_v t_0)$ the bilayer frame area is updated and with probability of $1 - P$, other trial moves are performed. To update the projected area: first, the box sides will be updated to $Lx \rightarrow Lx + \Delta Lx$ and $Ly \rightarrow Ly + \Delta Ly$ in which $\Delta Lx$ is chosen randomly in the interval $[\ 0.04 d_{dts}, 0.04 d_{dts}]$ and $\Delta Ly = \Delta Lx(Lx/Ly)$. The $X, Y$ coordinates at each vertex are then updated to $X \rightarrow (1 + \Delta Lx/Lx)X$ and $Y \rightarrow (1 + \Delta Ly/Ly)Y$. To accept or reject the move, Metropolis algorithm with the acceptance probability ($P_{acc}$) at temperature $T$ is performed, where:

$$P_{acc} = \min\left[1, \left[\frac{A_{p-new}}{A_{p-old}}\right]^{N_v} \exp\left(\tau \Delta A_p - \Delta E\right)\right] \qquad (2.1)$$

Note: We have set $k_B T = 1$, therefore all the energy unit in FreeDTS is in $k_B T$.

### Section 3: Osmotic pressure

To model the effect of osmotic pressure, FreeDTS can also apply an algorithm based on the Jacobus van 't Hoff equation, $\Pi = icRT$, where $i$ is van 't Hoff index, $c$ is the molar concentration of solute, $R$ is the ideal gas constant, and $T$ is the temperature. For a vesicle system, we define, $V_{ini}$ as the initial volume of the vesicle and effective compartment concertation of solute as $\bar{c}_j = \sum_n i_n c_n$, where $j$ is either inside (at $V_{ini}$), or outside and the summation run over all the solute types in the $j$ compartment. Therefore, we have

$$\Delta\Pi = RT\left(\frac{V_{ini}\bar{c}_{in}}{V} - \bar{c}_{out}\right) \qquad (3.1)$$

Energy associated with changes of the vesicle volume from $V_{ini}$ to $V$ will be obtained as

$$\Delta E_{osmos}(V) = -RT\int_{V_{ini}}^{V} \Delta\Pi dV = -RT\left[\bar{c}_{in}V_{ini}\ln\frac{V}{V_{ini}} - \bar{c}_{out}(V - V_{ini})\right] \qquad (3.2)$$

FreeDTS can directly use equation 3 to apply osmotic pressure difference. However, in most cases, this will be equivalent to the equation 3 in the main manuscript since

$$\Delta E_{osmos}(V) = RT\bar{c}_{out}V_{\text{eq}}\left\{\frac{1}{2}\left(\frac{V - V_{\text{eq}}}{V_{\text{eq}}}\right)^2 + \left[\sum_{i=3}^{\infty}\left(\frac{V_{\text{eq}} - V}{iV_{\text{eq}}}\right)^i\right]\right\} \qquad (3.3)$$

Where $\bar{c}_{out}V_{\text{eq}} = \bar{c}_{in}V_{ini}$ and $K = RT\bar{c}_{out}V_{eq}$. In typical experiments, $\bar{c}_{out}$ and $\bar{c}_{in}$ are in the order of $\sim 10mM$ and vesicle volume is $V \sim \mu m^3$, making, $K \sim 10^5 - 10^6 kT$ that is much larger than bending energy associated with vesicle deformations at this scale ($4\pi\kappa \sim 10^2 - 10^3\ kT$). Therefore, only the first term of equation 4 will be relevant and the vesicles will adopt to $V = V_{\text{eq}} = \frac{\bar{c}_{in}V_{ini}}{\bar{c}_{out}}$.

### Section 4: System evolution

System evolution and the equilibrium properties of the membranes are evaluated by standard Monte Carlo sampling of Boltzmann's probability distribution. We have set $k_B T = 1$, therefore all the energy unit in FreeDTS is in $k_B T$. Every Monte Carlo move consists of,

1) $N_v$ vertex positional updates.
2) $N_e$ Alexander moves: a trial to flip the mutual link between two neighboring triangles.
3) $N_i$ inclusion moves where $N_i$ is the number of the inclusions in the system. An inclusion move can be either rotation or a Kawasaki move.

In FreeDTS there are two options to perform these moves.

1) $N_v + N_e + N_i$ sweep attempts is one step. At each attempt, with probability of $N_v/(N_v + N_e + N_i)$ the selected move is vertex position update; with probability of $N_e/(N_v + N_e + N_i)$ the selected move is Alexander moves and with probability of $N_i/(N_v + N_e + N_i)$ the selected move is inclusion moves.
2) In one step $N_v$ vertex update, $N_e$ Alexander moves and $N_i$ inclusion moves will be performed.

If a system is coupled to the tension controlling scheme, then there will be also one trial move to change the box size with the a given probability (See section SI-2). There is also a possibility to activate parallel tempering algorithm. This will run multiple replicas at different temperature using OpenMP parallelization and after certain steps (defined by the user) it exchange the conformation of the replicas.

### Section 5: Benchmark: performance of the code for the Monte Carlo moves

In order to test the performance of the code, we measured the time elapsed for each move on four different systems. These systems are chosen to consist of the most (computationally) simple system (system 1, a closed triangulated surface) up to the most complex system (system 4: a periodic flat membrane fully decorated with the most complicated inclusions). The results were collected on a ThinkStation P620, model name AMD Ryzen Threadripper PRO 3995WX 64-Cores. Note, while the system contains 64 cores, the simulations were performed on a single core with a core frequency of "CPU max GHz 4.308 and CPU min GHz 2.200". In Table 2, the time elapsed between these moves is shown (reported for 1000 moves). As the number of calculations and, therefore, the duration of each move is dependent upon the results of the move (accepted or rejected and what is the cause of rejection), we monitored the time elapsed for each move in Figure SI-7-10. In these figures, only the uppermost lines are the accepted moves. below zero is a rejected move and the The simulation results presented in the main manuscript was obtained by simulations of $10^6 - 10^7$ steps which is resulting in 1-10 days of simulations on the described workstation. Please note a set of $N_v + N_e + N_i$ sweep attempts is called one MC step (see section SI-4).

| | Time Per 1000 Moves [millisecond] |
|---|---|
| Link Flip Move (Alexander Move) | 0.45-0.95 |
| Vertex Position Update Move | 24-32 |
| Inclusion Move | 1-1.1 |
| Box size update Move | 2.5 (per vertex) |

**Table 5.1: The range of expected time elapse for each Monte Carlo move.**

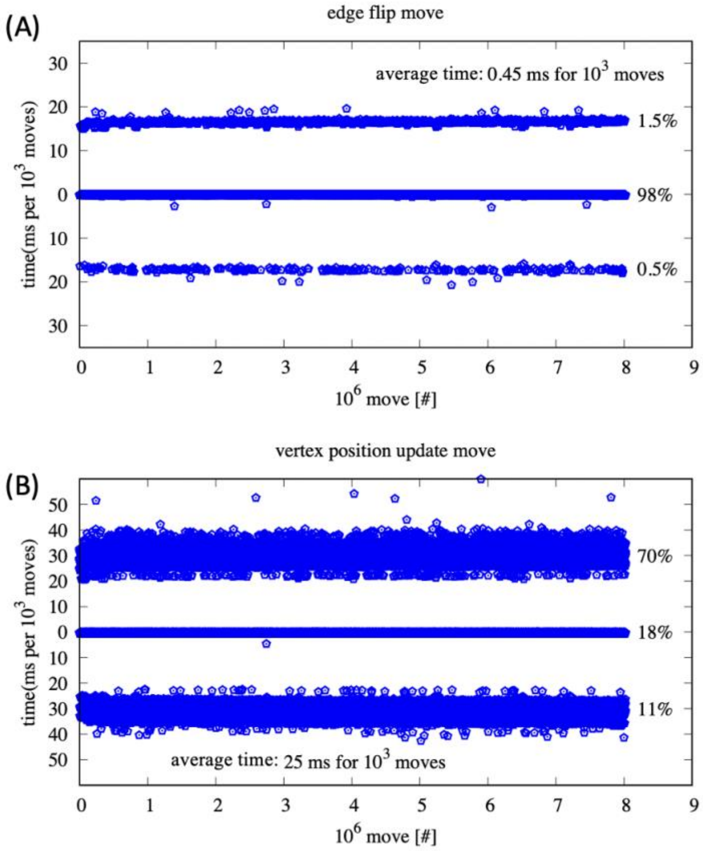

**Figure SI-5.1: A spherical triangulated mesh containing 802 vertices.** Only the uppermost line are the accepted moves. (A) Link flip attempts. (B) vertex position update.

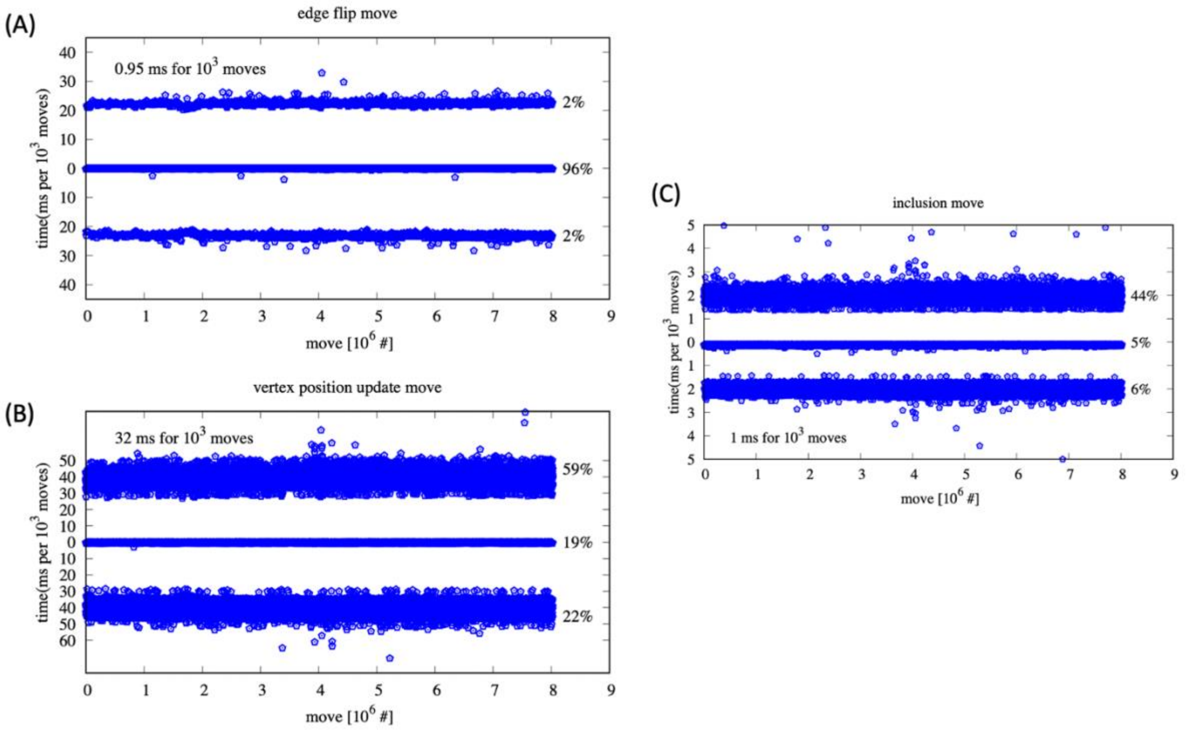

**Figure SI5.2: A spherical triangulated mesh containing 802 vertices and fully covered by inclusions.** Only the uppermost line are the accepted moves. (A) Link flip attempts. (B) vertex position update. (C) inclusion moves.

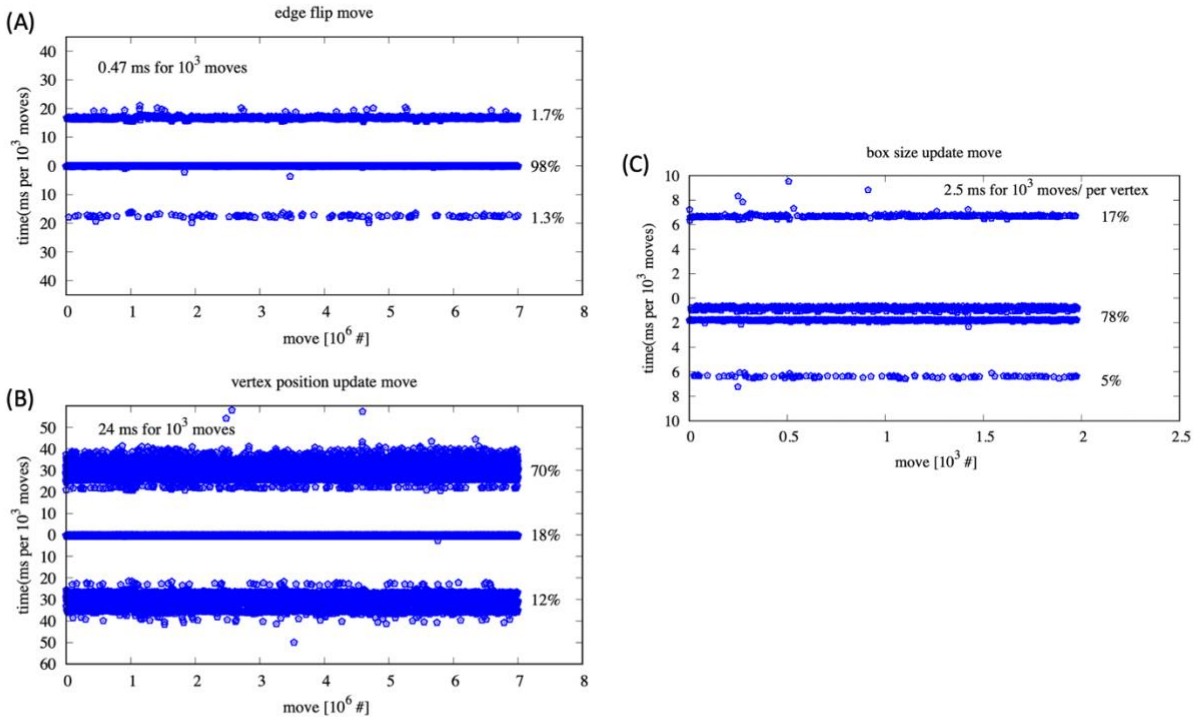

**Figure SI-5.3: A flat period triangulated mesh containing 700 vertices coupled with the tension controlling algorithm.** Only the uppermost line are the accepted moves. (A) Link flip attempts. (B) vertex position update. (C) box size update move.

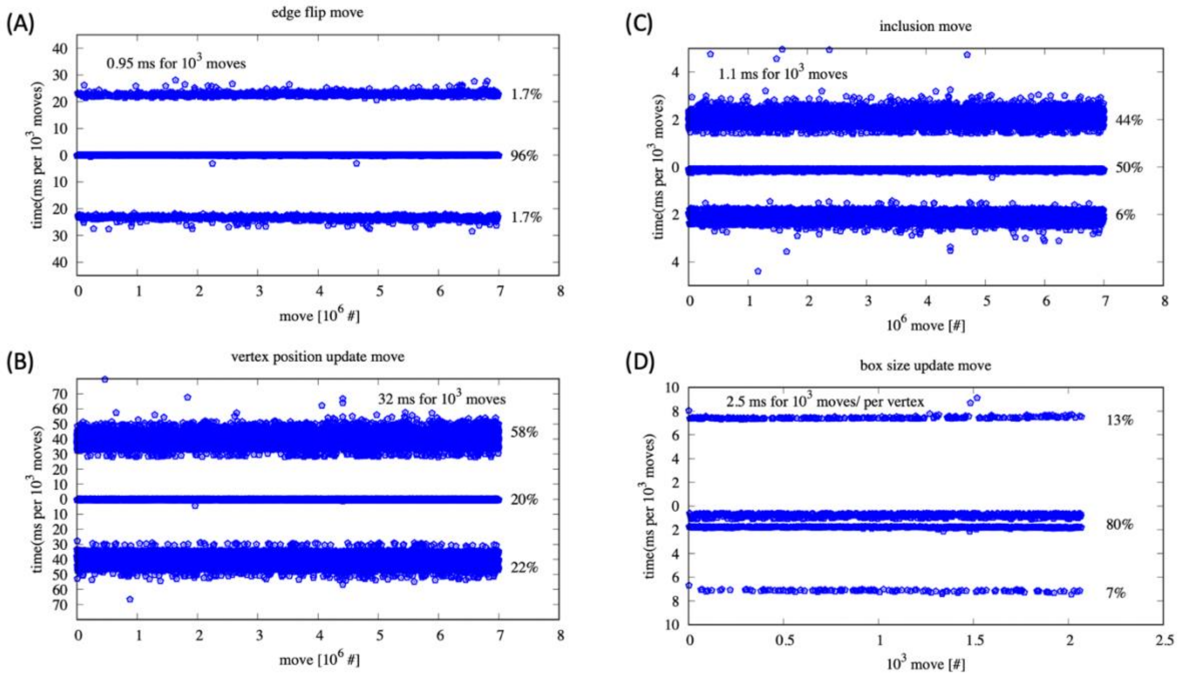

**Figure SI-5.4: A flat period triangulated mesh containing 700 vertices coupled with the tension controlling algorithm.** Only the uppermost line are the accepted moves. (A) Link flip attempts. (B) vertex position update move. (C) inclusion moves. (D) box size update move.

### Section 6: Converting $\boldsymbol{l_{dts}}$ to a physical unit

While it is possible and very common to explore membrane behaviors in terms of relative length scales (in the unit of $l_{dts}$) [1, 2] often it is more appealing to convert this length unit ($l_{dts}$) to a physical unit in particular if we are focusing on a specific real system. Previously we have done this for Shiga toxin, Annexin 4 and simple vesicles [3, 4]. However, this conversion scheme is not unique and can be adopted according to the phenomenon we are interested in. We briefly discuss this topic below and refer interested readers to the original articles [3-6].

**For system containing proteins:** To convert $l_{dts}$ to a physical unit e.g., nm, we assume that the area of a vertex which is $\sqrt{3}/2l_{dts}^2$ is equal to the area is equal to the area of the smallest circle wrapping the protein ($\pi R^2$) [3, 6]. Therefore,

$$l_{dts} \sim 1.9\,R \qquad (6.1)$$

***For pure bilayer:*** When two faces of a membrane approach each other, within nm distance, they repel each other through hydration and membrane protrusion forces that are below the DTS resolution. We account for these forces by treating them as infinite repulsion by restricting different segments of a membrane surface from approaching a distance at which these forces are effective (0.7 nm). Thus, $l_{dts}$ is the bilayer thickness plus a distance in which hydration and membrane protrusion forces are effective ($\sim 0.7nm$), leading to $l_{dts} \sim 4.5nm$. Note, for membranes with small deformations, $l_{dts} > 4.5nm$ nm can also be chosen. However, for highly deformed membranes, this choice implies that some realistic configurations of the membrane have been eliminated[4].

### Section 7: Extra Tables

| System | Area per vertex |
|---|---|
| Vesicle (i) | $1.545 \pm 0.008$ |
| Tensionless flat membrane (ii) | $1.548 \pm 0.007$ |
| Flat membrane with constant AP (iii) | $1.540 \pm 0.003$ |
| Tensionless flat membrane with inclusions (iv) | $1.547 \pm 0.007$ |
| Tensionless flat membrane between two walls (v) | $1.546 \pm 0.006$ |

**Table SI-7.1: Area per vertex remains constant for membrane simulations in different conditions. Also see Figure 1SI.**

### Section 8: Extra Figures

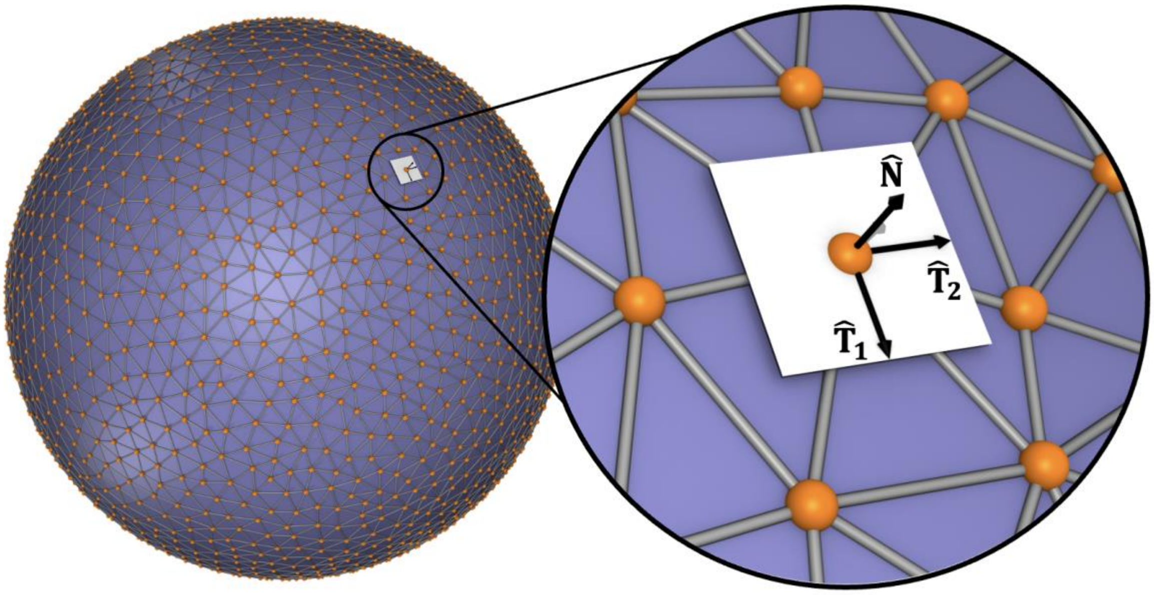

**Figure SI-8.1:** A regular triangulated surface containing $N_v$ vertices, $N_e$ edges and $N_T$ triangles. Using a set of discrete gematrical operations we obtain, principal curvatures $(c_1, c_2)$, and principal directions $(\hat{\mathbf{T}}_1, \hat{\mathbf{T}}_2)$, and a surface normal $(\hat{\mathbf{N}})$ on each vertex.

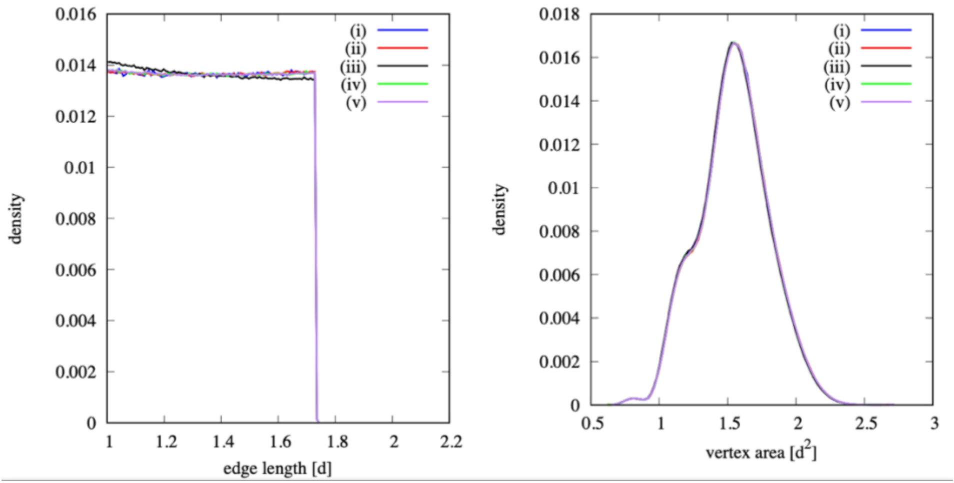

**Figure SI-8.2: Distributions for the systems shown in the Table 1SI.**

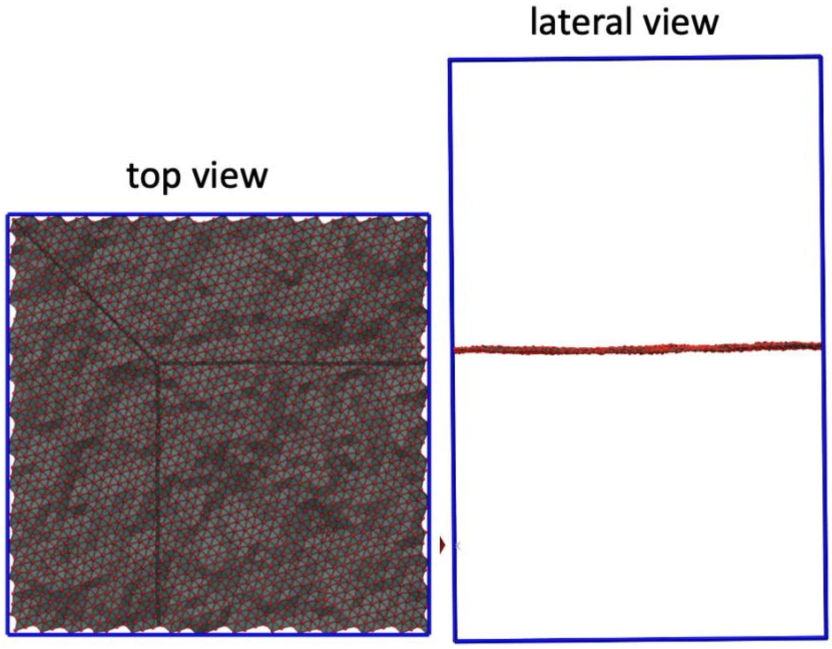

**Figure SI-8.3: Two different views of a triangulated surface in a periodic box. Note, the vertices along each edge are connected with those on the opposite side through the periodic box. This connectivity cannot be visualized.**

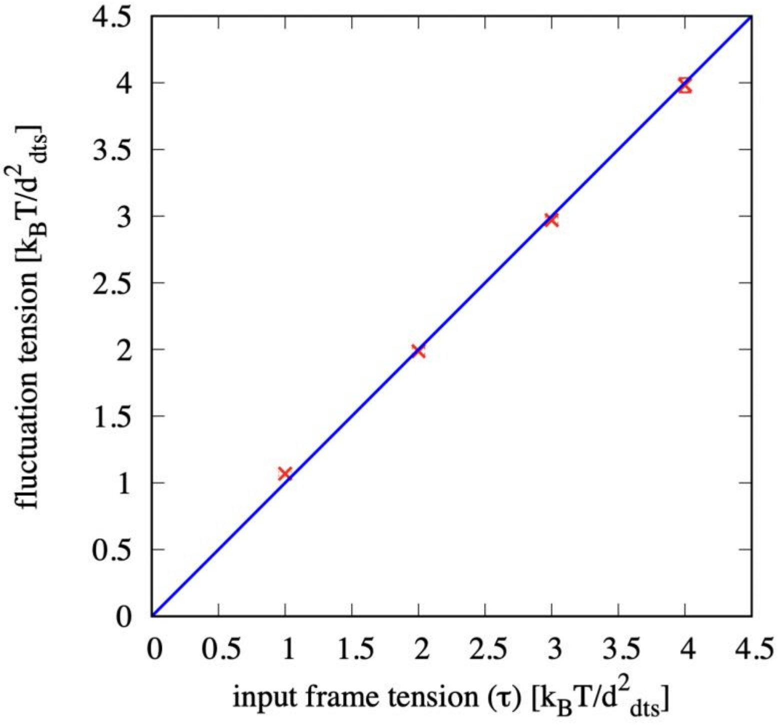

**Figure SI-8.4: The membrane tension determined by the undulation spectrum as a function of the input frame tension. These results indicate that they are equal.**

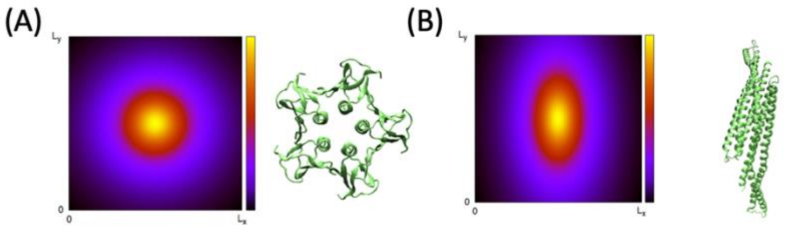

**Figure SI-8.5: Two different protein types in FreeDTS.** (A) Protein type 1, in which its interaction with the membrane modifies the membrane in a symmetric (or almost symmetric) way (B) Protein type 1, in which its interaction with the membrane modifies the membrane in a asymmetric way.

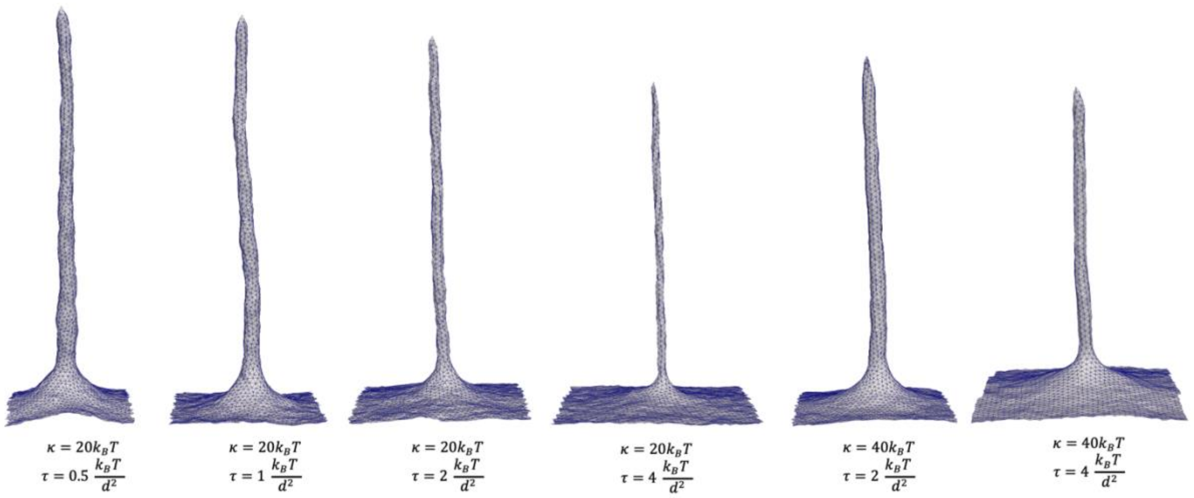

**Figure SI-8.6: Tube pulling with different size.** By changing membrane tension or bending rigidity, tubes with different radii will form.

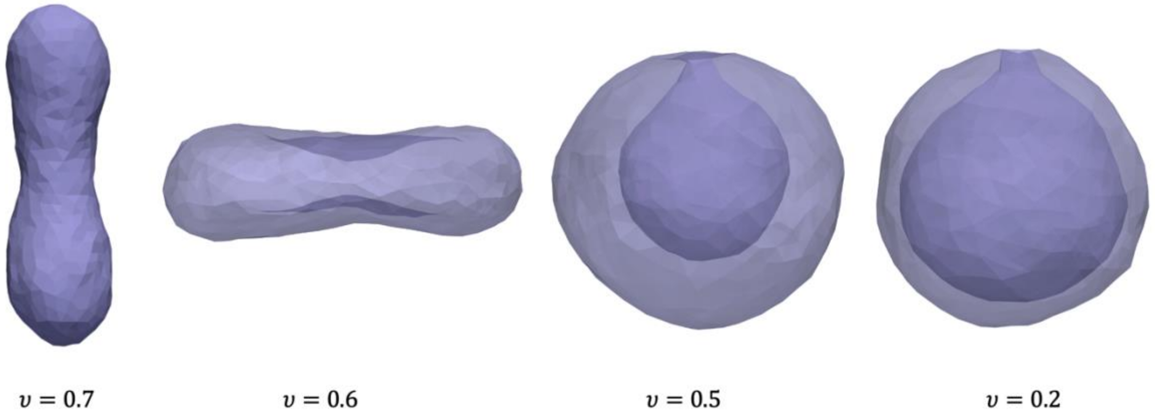

**Figure SI-8.7:** Transition of a spherical vesicle from prolate-to-oblate and oblate-to-stomatocyte by volume reduction.

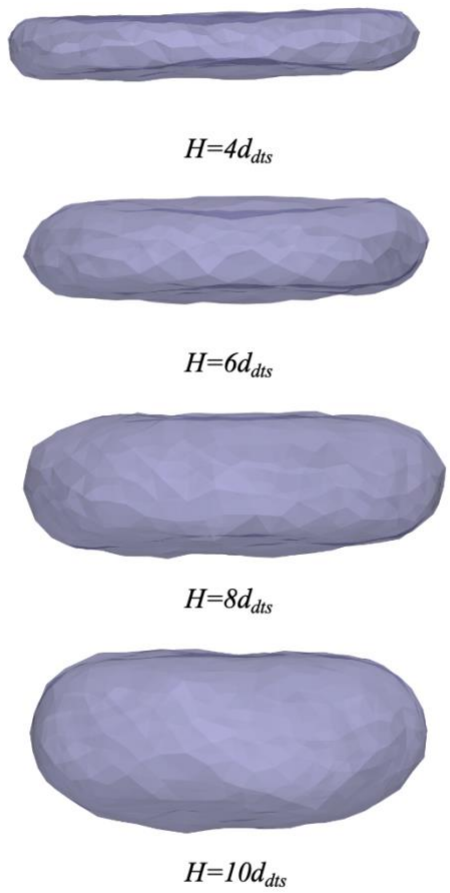

**Figure SI-8.8:** A vesicle sandwiched between two confining walls in the Z direction for different distance between the walls.
